# Early death and neuronal abnormalities in *depdc5* loss-of-function mosaic zebrafish models

**DOI:** 10.64898/2025.12.04.692362

**Authors:** Sneham Tiwari, Christopher M. LaCoursiere, Hyun Yong Koh, Joseph Pascucci, Parul Chaudhary, Richard D. Goldstein, Annapurna Poduri

## Abstract

*DEPDC5* (DEP domain-containing protein 5) encodes a repressor of the mTORC1 signaling pathway. Variants in *DEPDC5* are associated with a range of focal epilepsies, including mosaic variants associated with focal cortical dysplasia (FCD) and other focal brain malformations with brain-only somatic mosaic variants. To investigate the role of *DEPDC5* in human epilepsy related to mosaic variants, we have generated mosaic *depdc5* loss-of-function zebrafish models using homology-based constructs acutely targeting *depdc5* and labeled with tdTomato to allow for visualization of the degree of mosaicism. The resulting mosaic *depdc5* CRISPants demonstrated early larval death, with ∼50% of CRISPants (vs. 10% of controls) dead by 7 days post fertilization (dpf), analogous to the early death sometimes associated with human *DEPDC5*-related epilepsy. We compared *depdc5* CRISPants with uninjected and scrambled controls from the same clutches. Body and head size were reduced in the *depdc5* CRISPants. Analysis of swimming behavior showed a striking reduction in distance traveled and maximum velocity in the *depdc5* CRISPants vs. controls. Based on visual confirmation of mutational load, we categorized CRISPants into *depdc5*+ vs. depdc5++, reflecting weak vs. strong tdTomato fluorescence. We observed that *depdc5*++ CRISPants had increased episodes of posture loss, suggesting increased seizure-like behavior related to higher percentages of mutant cells. Local field potential recordings revealed increased neuronal hyperexcitability in *depdc5* CRISPants vs. controls. Acridine orange staining demonstrated early apoptosis in the CRISPants vs. controls. Our mosaic *depdc5* CRISPants provide a clinically relevant model to study the role of mosaic DEPDC5-related epilepsy and early death.

## Introduction

Epilepsy is a common, life-affecting disease with burgeoning genetic discovery in critical need of functional studies.^1–3^ It has a lifetime prevalence of 1 in 26.^4, 5^ At least one-third of individuals with epilepsy do not achieve seizure control with anti-seizure medications,^6^ and often experience neurodevelopmental comorbidities, such as intellectual disability, autism spectrum disorder, and psychiatric disorders.^7, 8^ Studying tissue from human epilepsy surgery specimens, our group and others have implicated *de novo* post-zygotic mutation^9^ resulting in cellular mosaicism as one of the etiologies of focal epilepsy, with genetic variants isolated to one cerebral hemisphere or one region of brain.^10–20^ In epilepsies with or without structural lesions on neuroimaging, mosaic variants are present in a subset of brain cells, many affecting genes in the mammalian target of rapamycin^5^ pathway, e.g., *DEPDC5, AKT3*, *PIK3CA*, and *MTOR* itself.^10–19, 21, 22^

The mTOR pathway is important for several aspects of intracellular function, including the regulation of cellular growth and cell proliferation.^17, 21, 23–32^ In the brain, the mTOR pathway is active early during development and is implicated in neuronal differentiation and growth, synaptogenesis, and dendrite formation, thus playing a key role in the structure and function of the cerebral cortex^33^. The gene *DEPDC5* encodes a GTPase that, together with *NPRL2* and *NPRL3* (Nitrogen Permease Regulator-like 2 and 3), constitutes the GATOR-1 (gap activity towards RAGS-1) complex, an upstream inhibitor of mTOR activity.^34, 35^ ^36–39^ Mutations in *DEPDC5* cause a broad spectrum of focal epilepsies.^21, 27, 32, 36, 37^ *DEPDC5* variants were initially described in familial epilepsies with autosomal dominant inheritance^36, 37, 40^ and sporadic epilepsies, including cortical malformations.^41, 42^ *DEPDC5*-associated disease-causing germline variants are predominantly heterozygous and inactivating, including nonsense, frameshift and splice-site variants, with missense variants presumed to result in loss of function. Haploinsufficiency of DEPDC5 leads to decreased mTOR inhibition by the GATOR-1 complex, leading to cellular abnormalities and disrupted cortical lamination (FCD),^21, 23, 43, 44^ neuronal hyperexcitability, and seizure susceptibility. ^20, 37, 45, 46, 21, 40^ When *DEPDC5* variants have been identified in focal epilepsy resections, they have been present with less than 50% variant allele fraction in brain tissue and absent in other tested tissues, indicating the presence of a somatic mutation, or a mosaic variant that arose post-zygotically in the developing brain.^13, 14, 16, 21, 47^ Additionally, for focal lesions in the presence of a germline loss-of-function *DEPDC5* variant, a two-hit model has been proposed that involves a second, somatic, loss-of-function variant.^5^ Compound heterozygous variants (germline plus somatic) in *DEPDC5* have been reported, supporting this model.^48–54^ Additionally, our research on sudden unexpected deaths in the pediatric age range (SUDP) identified variants in *DEPDC5* in 2 infants who died at 3 months and 5 months of age, raising questions about the role of seizures and early death.^55^

Vertebrate animal models have played a critical role in validating the role of *DEPDC5* in epilepsy via altered mTOR activity.^56–59^ *Depdc5* knockout zebrafish models have shown spontaneous seizure-like activity, increased neuronal excitability, and sensitivity to convulsants.^48, 51, 57^ Further, neuronal-specific conditional *Depdc5* knockout mice displayed mTORC1 hyperactivation and, rarely, exhibit seizure-related mortality in adulthood, whereas constitutive *depdc5* knockout results in embryonic lethality as well as evidence of mTOR hyperactivation (increased phosphorylation of S6).^58–60^

To understand mosaic *DEPDC5*-related human epilepsy and associated phenotypes, we have utilized the larval zebrafish (*Danio rerio*) system to study seizure-related phenotypes that result from loss of *depdc5* function. CRISPR/Cas9 gene editing has the advantage that its efficiency is typically incomplete, such that introducing guides and Cas9 mRNA or protein at the 1-cell zygote results in mosaic organisms in most cases. Targeting *depdc5* in zebrafish larvae, we address the pathogenesis of mosaic epilepsy in this clinically relevant experimental animal model system. Here we demonstrate that *depdc5* larval mosaic models recapitulate features of human epilepsy caused by mosaic *DEPDC5* variants, including seizures, hyperexcitability, and early death.

## METHODS

### Zebrafish welfare and husbandry

All zebrafish experiments were approved and conducted in accordance with the standards of the Boston Children’s Hospital (BCH) Institutional Animal Care and Use Committee (IACUC). Zebrafish were maintained in the Aquatic Resources Program (ARP) at BCH, at 28.5°C^61^ with 12-hour dark-light cycles in a dedicated zebrafish facility. Zebrafish larvae were raised in sterile fish water with 0.01 mg/L methylene blue and moved to sterile fish water on 1 day post fertilization (dpf). All experiments were performed in the *Danio rerio* Casper strain.

### Generation of *depdc5* mosaic zebrafish embryos

To generate *depdc5* loss-of-function models, we developed a homology-based modular construct with complementary DNA, splicing acceptor, and Gal4-VP16–UAS-driven tdTomato reporter system (Figure 1), modified from a study that utilized microhomology-mediated gene editing in zebrafish.^62^ To target the zebrafish *depdc5* gene (sole ortholog of human *DEPDC5)*, we used CHOPCHOP^2, 63^ to design a highly specific guide RNA (gRNA) targeting the intron between exons 9 and 10 (tggttgcttagcagtgttggggg) to generate a specific premature stop codon. Briefly, we designed 48bp upwards and downward homologous arm (DHA) oligonucleotides, and the UHA (tttggcttaactgttgtcaaaaagcaaggcatggttgcttagcagtgt) and ligated then with a linearized UFlip vector using standard techniques^62^.

**Figure 1.**
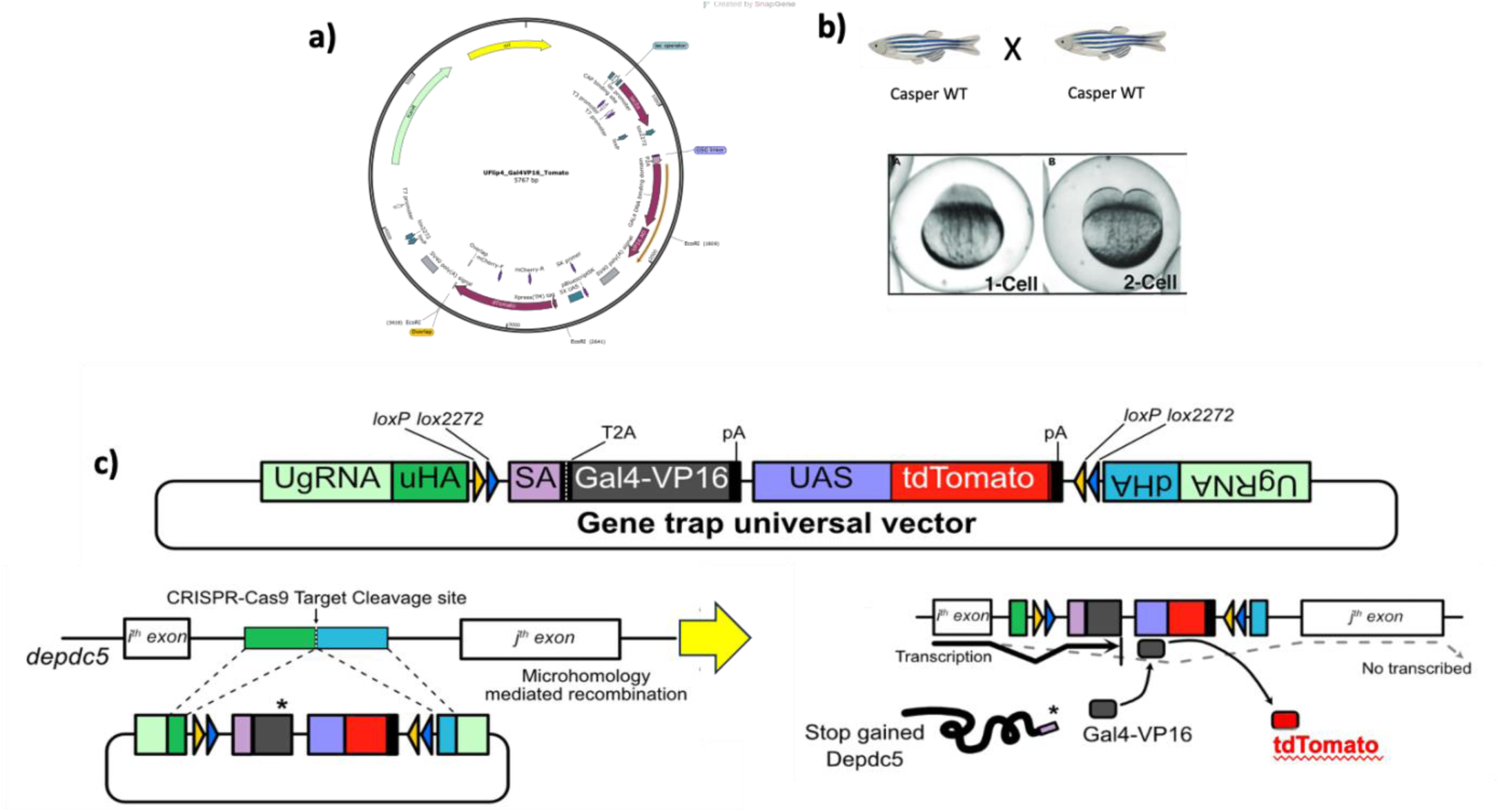
Figure 1. Generation of readily identifiable mosaic mutant *depdcS* zebrafish embryos. a) Plasmid construct depicting [uHA, dHA, Gal4NP16/UAS and tdTomato] b) We breed *Casper* WT fish to generate transparent embryos, into which constructs can be injected at the 1-cell or 2-cell stage. c) The construct strategy displaying the cut by CRISPR-Cas9.CRISPR/Cas9 enzyme activity leads to a stop gained during *depdc5* transcription, activating tdTomato fluorescence , which is utilized to sort *depdc5* CRISPants (fluorescent) vs. mutation- negatives (non-fluorescent) embryos.

Using established microinjection protocols^3, 64^, we performed microinjections utilizing the construct, *depdc5* gRNA and CRISPR/Cas9 mRNA. Concentrations were titrated to achieve low to intermediate rates of mosaicism (variant allele fraction), with the optimal concentrations determined to be 25pg *depdc5* gRNA, 25pg UgRNA, 10pg construct, and 300pg *Cas9* mRNA in a 2nl/embryo final concentration. Scrambled gRNA (acaaggaggtaggcgagaac)^3^ was utilized to generate handling/experimental control embryos. Microinjections were delivered into the one cell of the 2-cell stage to ensure mosaicism. Fluorescence conferred by the tdTomato marker gene was evaluated as an indicator of successful insertion of the *depdc5*-targeting construct; embryos were sorted based on red fluorescence at 1 dpf from the negative (non-fluorescent) embryos. For the first clutch of embryos, we used Sanger sequencing to confirm that the injected embryos demonstrated evidence of successful CRISPR/Cas9 gene editing (forward primer aaagcattacaagcaacatgagt and reverse primer tgggttttcattgtgtaaggtgt). For subsequent experiments, we sorted embryos based on visual red fluorescence to identify those carrying the mutation. In our F_0_ microinjected CRISPants, we assessed the efficiency of CRISPR/Cas9 gene editing and resulting degree of mosaicism using visual confirmation of red fluorescence intensity. Based on relative fluorescence intensity at 1 dpf, larvae were classified as *depdc5*+ reflecting lower fluorescence intensity (<30% brightness) vs. *depdc5++* reflecting higher fluorescence intensity (100% brightness).

### Survival analysis

Casper wildtype (WT) zebrafish were mated to generate fertilized oocytes (zygotes), and the resulting embryos were microinjected at the 2-cell stage with depdc5-targeting gRNA, construct, and Cas9 mRNA. Embryos were seeded on 10cm^2^ transparent petri plates (30 embryos/plate) and sorted based on red fluorescence at 1 dpf, as above. Red fluorescent-positive *depdc5* CRISPants were observed for 14 days. Sterile water was refreshed every day, and embryos were fed with live rotifer solution daily after 6 dpf. *Depdc5* CRISPants were counted and assessed for larval death. Dead larvae were removed from the petri plates, and ‘missing’ larvae were presumed to have died and dissolved. Uninjected controls from the same clutch and *scrambled* controls were similarly counted. Results were graphed and analyzed using Kaplan-Meier survival analysis.

### Larval head and body measurements

Larval head size and body length were determined at 4 dpf by mounting the larvae in 1.5% low-melting agarose (Fisher Scientific, Cambridge, MA, USA), dorsal side up. Head size was measured as the linear distance between the right and left lens^65^, and body length was measured as the linear distance from snout to the end of the tail excluding the fins. For malformed CRISPants, Image J (Fiji) software curve line tool was used to measure the length of the animal. Each value was noted and then compared to uninjected and *scrambled* controls from the same clutch. Measurements and calculations were determined with the ImageJ 1.41(NIH, USA) for each embryo.

### Swimming behavioral assay and analysis

5 dpf larvae were placed in clear 96-well plates, 1 larva/well in 100ul of sterile fish water. Using the Noldus DanioVision visualization and recording system (Leesburg, VA) spontaneous swimming parameters were recorded for 30 minutes in the dark, as previously described^2, 3^. Larvae were monitored for normal vs. seizure-like swimming activity. We graphed the average distance travelled from the center of the well and the maximum velocity using Prism 10 analysis software, to compare *depdc5* CRISPants vs. the uninjected and *scrambled* control larvae.

Further, for advanced visualization and tracking, we utilized the Kestrel multi-camera array microscope (MCAM, Ramona Optics Inc., Durham, NC, USA) to analyze spontaneous swimming behavior and behavior following administration of the proconvulsant pentylenetetrazole (PTZ) (5mM) in 5-minute trials in the dark. 5 dpf *depdc5* CRISPants, uninjected controls, and *scrambled* control larvae were placed in clear 96-well plates, 1 larva/well, in 100ul of sterile fish water. After a 10-minute buffer time in the wells, the plate was loaded onto the Kestrel multi-camera array microscope (MCAM, Ramona Optics Inc., Durham, NC, USA). Baseline and PTZ readings were taken according to a protocol adapted from the Kestrel handbook and a recent study.^66^ Spontaneous baseline readings were acquired for 5 mins, then 50ul of sterile water was removed from the well and replaced with 50ul of 10mM PTZ to a final concentration of 5mM PTZ in 100ul sterile fish water/well. Five minutes later, the PTZ-induced Timepoint 1 readings were acquired for 5 mins. Tracking analysis was run on these trials and the generated metadata raw files were analyzed by modifying an analysis script published in a recent study.^66^ The values per frame were then consolidated for each larva and assessed for larval behavior ontologies, defined as i) Stationary, ii) Normal Swim, iii) Whirlpool, iv) Convulsion, and, v) Posture Loss (lateral orientation).^67^

### Local field potential recording and analysis

Electrophysiological recordings were obtained from larval zebrafish using established methods to record local field potentials (LFPs)^2, 3^. Larval zebrafish (5-6dpf) were paralyzed with alpha-bungarotoxin (2 mg/ml) (Invitrogen, Waltham, USA)^68^ and embedded dorsal side up in 50μl of 1.2% low-melting point agarose on a recording chamber. The chamber was placed on a Nikon Eclipse FN1 microscope. A borosilicate glass electrode (1-7 MΩ_resistance) filled with 2 M NaCl was placed in the optic tectum using a micromanipulator with visual guidance under the light microscope. Clampex software (Molecular Devices, Sunnyvale, USA) was used for the LFP recordings, which were performed in current-clamp mode, by standardized protocol in the lab^3^. After a baseline recording of 30-minutes, 40 mM PTZ (dissolved in the recording solution) was added to the recording chamber to confirm the correct placement of the probe in the tectum, as classic PTZ specific large spikes were observed in the reading. We continuously observed larvae for heartbeat and blood circulation in the head and included recordings only from larvae maintaining a visible heartbeat. We excluded from the analyses larvae that died during exposure to alpha-bungarotoxin or during the 30-minute experimental trial.

All electrophysiology recordings were analyzed, blinded to genotype, using a custom MATLAB software framework with the established electrophysiology protocol in the lab^3^. Briefly, the framework detects the noise floor (0.006–0.012 mV) that aids in detecting the spikes adjusted to ambient activity. Larvae were categorized into different groups as i) non-seizing (normal or quiet readings), or ii) seizing (with few high amplitude or frequent high amplitude spikes) referred to as small- vs. large-amplitude events^69^ based on signal amplitude. We further quantified the non-seizing and seizing larvae as per the two defined categorized groups and quantified the proportion. The *depdc5* vs. scrambled and *depdc5* vs. control were analyzed using Chi-square (and Fisher’s exact) test and data were illustrated using Prism 10 analysis software.

### Assessment for apoptosis

Regarding the early death phenotype observed, we hypothesized that apoptosis at an early developmental stage might contribute. We performed Acridine orange (AO) (Sigma, USA) staining to visualize early apoptotic cell clusters. 4 dpf larvae were exposed to 5μg/ml AO in sterile fish water for 30 minutes, washed twice with sterile fish water, and sacrificed on ice. These larvae were embedded in 1.5% low-melting agarose, imaged using a Leica SP8 confocal microscope, and observed for green fluorescence. We counted apoptotic cell clusters in the larval tectum using the particle analysis function of ImageJ 1.41 (NIH, USA). All thresholds were set and used for the analysis of every image. Readings were analyzed and graphed using Prism 10.

### Western immunoblots (Supplementary data)

Embryos at 5 dpf from CRISPants and uninjected controls were homogenized in 25μl RIPA buffer and sonicated on ice. Homogenization was continued until a uniform lysate was obtained. Homogenates were centrifuged for 10 minutes at 4°C at 13000 rpm to pellet cellular debris, supernatant was retained, and protein was quantified using the BCA kit (Pierce BCA Protein Assay Kit, Thermo Scientific, Rockford, IL). Each protein sample was subjected to SDS-PAGE gel (12.0%) at a concentration of 18ug/25ul. Protein was dry transferred to an Invitrogen iBLOT2 PVDF membrane utilizing the iBLOT2 transfer station (Invitrogen, USA). The phospho-S6 (P-PS6) (Cell Signaling Technology, Danvers, MA) antibody was used (1:1,000) with LICOR secondary anti-rabbit antibody (1:20000). Beta-actin was used as a loading control, and the blots were imaged using a Biorad ChemiDoc MP Imaging System (USA).

### Statistical analyses

*Depdc5* CRISPants were compared to uninjected and *scrambled* controls. All statistical analyses were performed using GraphPad Prism 10. Values represent mean ± SEM. Differences among groups were analyzed by one-way ANOVA followed by Fisher’s LSD *post hoc* test or *t*-test. Comparison of survival among the experimental groups was performed using Kaplan-Meier survival analysis. For electrophysiological experiments, we compared the seizure vs. non-seizure events in i) *depdc5* CRISPants vs. clutch specific uninjected control and ii) *depdc5* CRISPants vs. scrambled controls, using the Chi-square and Fisher’s exact tests. A *p*-value of *<*0.05 was the threshold for determining statistical significance. All experiments were conducted with three technical replicates.

## Results

To understand the role of *DEPDC5* in neurodevelopmental epilepsy and related conditions, we developed a CRISPR/Cas9-mediated mosaic *depdc5* zebrafish model. We demonstrate morphological, behavioral, electrophysiological, and neuronal abnormalities in mosaic *depdc5* zebrafish larvae during early development and recapitulate the hyperexcitability characteristic of human epilepsy as well as the propensity to premature death.

### Generation and validation of the *depdc5* mosaic zebrafish models

To study the role of mosaic *depdc5* loss of function, which is associated with phenotypes in the zebrafish model system, we developed a homology-based modular construct with the Gal4-VP16–UAS-tdTomato reporter system and co-injected this construct with Cas9 mRNA and *depdc5* gRNA (Figure 1). We observed red fluorescence in larvae that had undergone successful Cas9-mediated gene editing, and distinguished *depdc5* CRISPants (fluorescent) (Figure 2c) from wildtype larvae (non-fluorescent) (Figure 2a,b) at 1 dpf. CRISPants were confirmed for a successful targeting of the *depdc5* gene by Sanger sequencing (Figure S1). Additionally, we demonstrated that *depdc5* modulation affects the mTORC1 pathway as indicated by an increase in the phospho-S6 protein, a downstream effector of the mTOR pathway,^70–72^ in *depdc5* CRISPants compared to uninjected controls from the same clutch (Figure S2).

**Figure 2.**
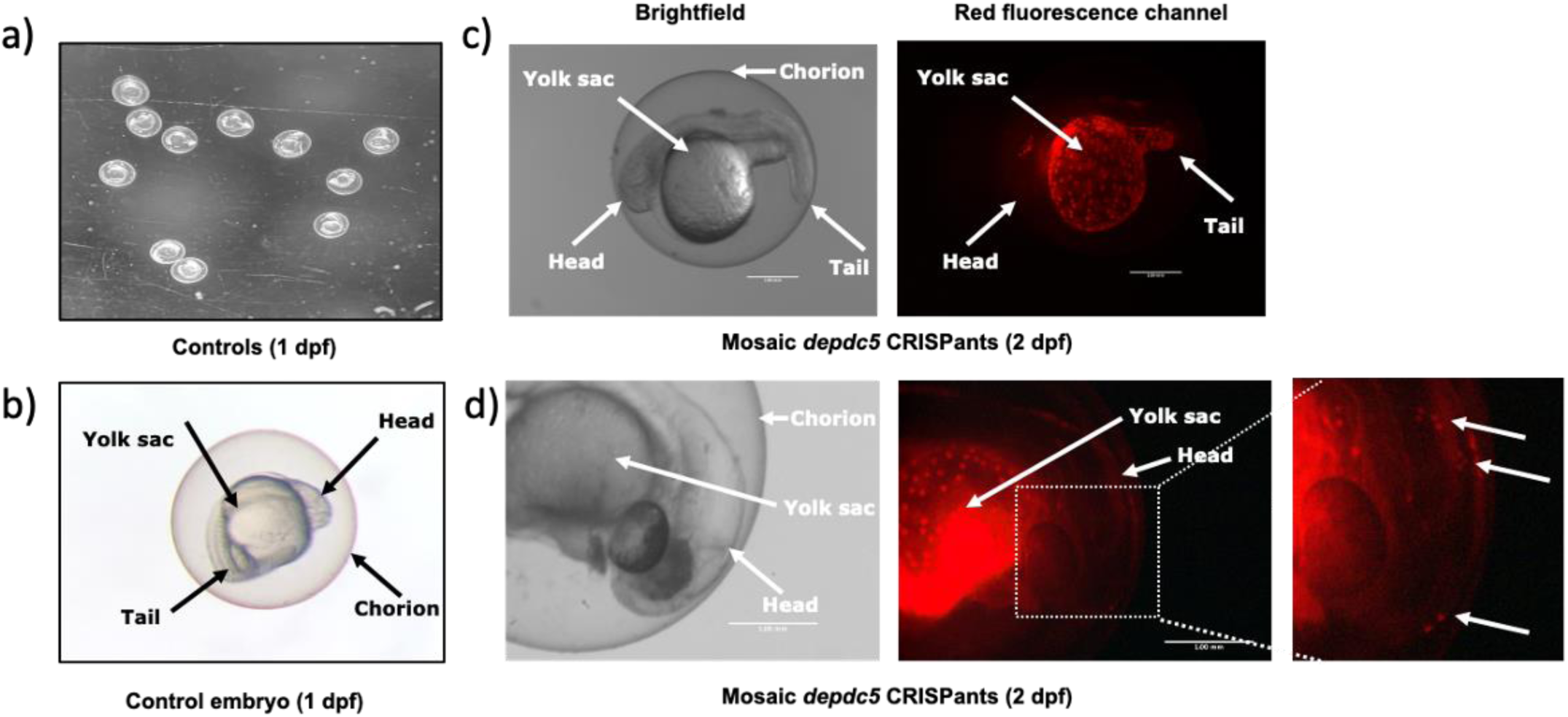
Visual confirmation of the mosaic mutations in the embryos. Under the red fluorescence, the microinjected embryos were visualized and subjected to fluorescence-based sorting into *depdc5* mosaic CRISPants. a) 1 dpf uninjected control (non- fluorescent) embryos in a petri-dish, b) representative embryo morphology of a control embryo (dpf1), c Brightfīeld and fluorescent *depdc5* CRISPants (fluorescent), showing mosaic mutations and d) *depdc5* CRISPant displaying mosaic mutations in the head region of the embryo (including inset showing focus on fluorescence in the head region) (Scale bar = 1mm)

To further assess the successfully generated CRISPants models for degree of mosaicism as well as the distribution of mutant cells in the mosaic larvae, including the head region, we assessed the brightness and the location of the mutant cells that were observed in the larvae (Figure 2c, 2d). We designated as *depdc5*++ larvae that were brightly fluorescent (100% brightness) whereas as any larvae with weak fluorescence (<30% brightness) were referred to as *depdc5*+ CRISPants.

### *Depdc5* mosaic CRISPants demonstrate early death

Since *DEPDC5* has been associated with sudden death in individuals with epilepsy (Sudden Unexpected Death in Epilepsy, or SUDEP) as well as Sudden Unexpected Death in Pediatrics even in the absence of epilepsy.^73–75^ To study the survival of *depdc5* CRISPants, we assessed survival in *depdc5* CRISPants vs. uninjected and *scrambled* controls. We observed death of *depdc5* mosaic CRISPants as early as 1-2 dpf. Overall, >50% of all *depdc5* CRISPants across multiple experiments were dead by 7 dpf, recapitulating the sudden death phenotype observed in *DEPDC5*-related SUDEP and SUDP, vs. only ∼10% in the uninjected (p<0.0001) and ∼15% in the *scrambled* control groups (Figure 3). This finding strongly indicates that *depdc5* loss of function is leading to early death.

**Figure 3.**
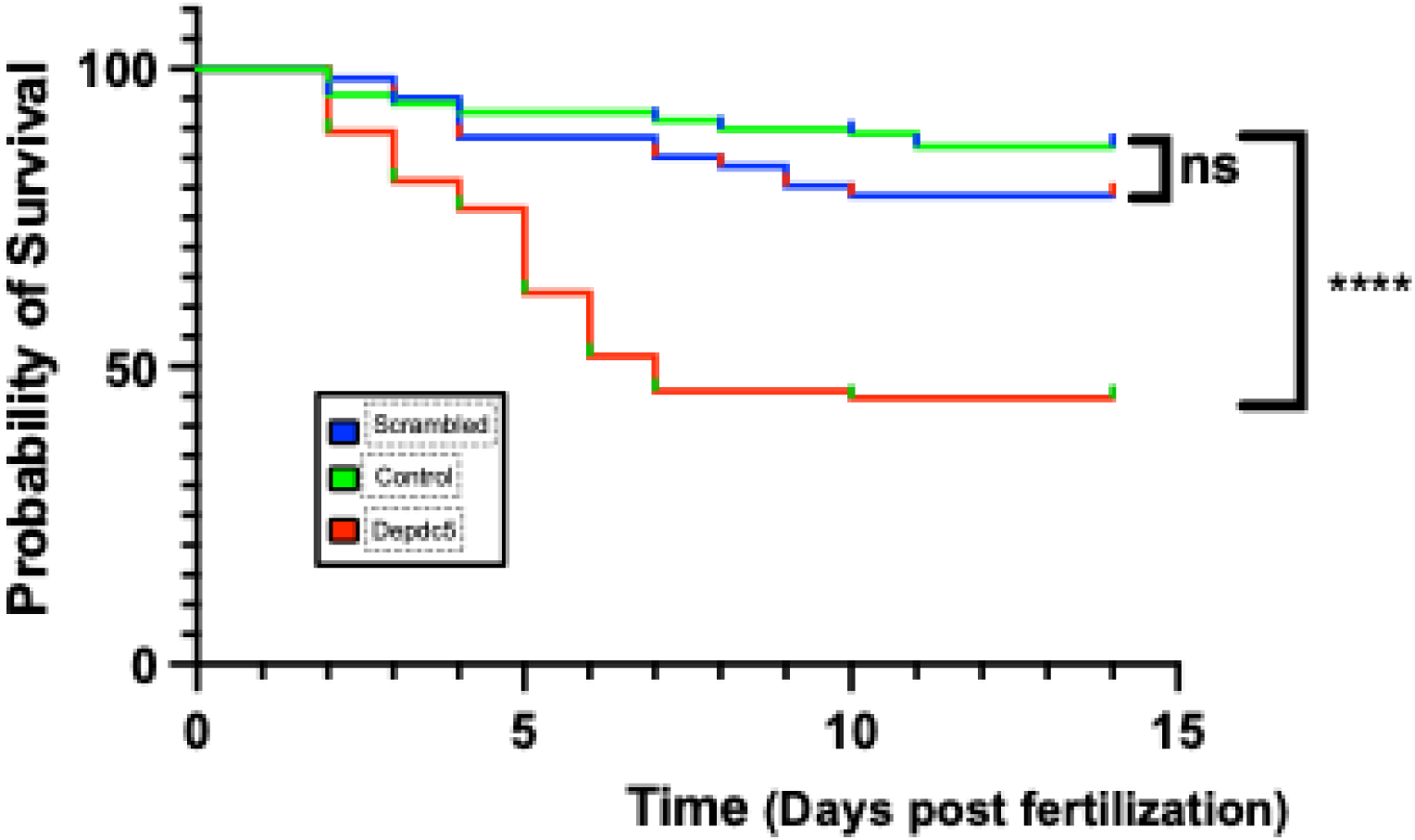
*Depdc5* CRISPants survival analysis. Kaplan-Meier demonstrating probability of survival (dpfθ*15). It describes the curve death starting at 1-2 dpf in mosaic *depdc5* CRISPants (red) and loss of ∼5O% CRISPants larvae by 7 dpf. The uninjected control (green) and scrambled larvae (blue) have a survival of -90%. control, n=138, scrambled, n=61, *depdc5* CRISPants, n=85, Kaplan-Meier Survival Comparison, p<0.0001

### Abnormal head and body sizes in *depdc5* mosaic CRISPants

We analyzed the head size and body length of 4 dpf *depdc5* CRISPants vs. uninjected controls and *scrambled* clutch mates. We observed that CRISPants had morphological changes compared to controls (Figure 4a), as well as pericardial and abdominal edema (data not shown). Importantly, *depdc5* CRISPants demonstrated a reduction in body length (Figure 4b, e) compared to uninjected and *scrambled* controls. We also observed that in comparison to controls’ straight body shapes, the CRISPants were malformed. Further, while we observed a reduction in the head size of *depdc5* CRISPants (Figure 4c, d), we also observed increased head-to-body ratios in *depdc5* CRISPants compared to controls (Figure 4f).

**Figure 4.**
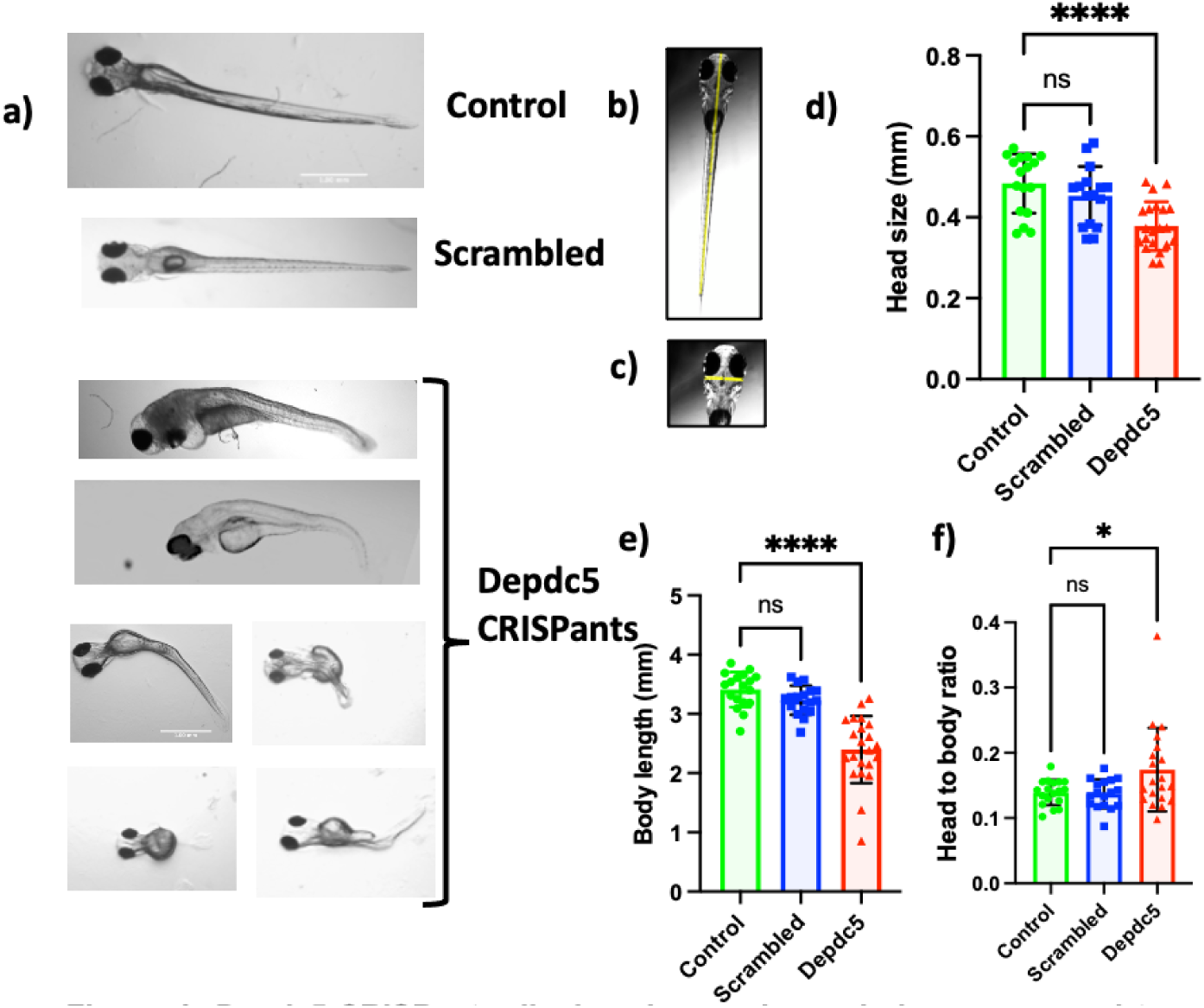
*Depdc5* CRISPants display abnormal morphology compared to uninjected and scrambled control. a) Abnormal morphology of the mosaic *depdc5* CRISPants compared to uninjected and scrambled control larvae b) representative larva to demonstrate measurement of body length (yellow line), and c) head size (yellow line), d) graphical representation of the head size, e) body lengths comparisons, and f) head to body ratio of *depdc5* CRISPants compared to clutch specific uninjected and scrambled controls, control, n=18; scrambled, n=17; *depdc5* CRISPants n=23 (Scale bar = 1mm) Ordinary One-way Anova test, (p<0.0001)

### Abnormal spontaneous and PTZ-induced seizure-like swimming patterns in *depdc5* CRISPants

In order to study whether *depdc5* loss of function increases the propensity to display seizures in *depdc5* CRISPants, we evaluated the swimming patterns of *depdc5* CRISPants vs. uninjected and *scrambled* controls using the Noldus DanioVision recording platform. First, we observed that *depdc5* CRISPants exhibited a 50% decline in the average distance travelled in a 30-minute observation period (Figure 5a, 5e) when compared to uninjected and *scrambled* controls. Proportionally, we observed a reduction in velocity of the *depdc5* CRISPants when compared to control larvae (Figure 5b). We attributed the wide range of velocities among *depdc5* CRISPants to their abnormal and variable morphologies, including bent spines and short body lengths (Figure 4a).

**Figure 5.**
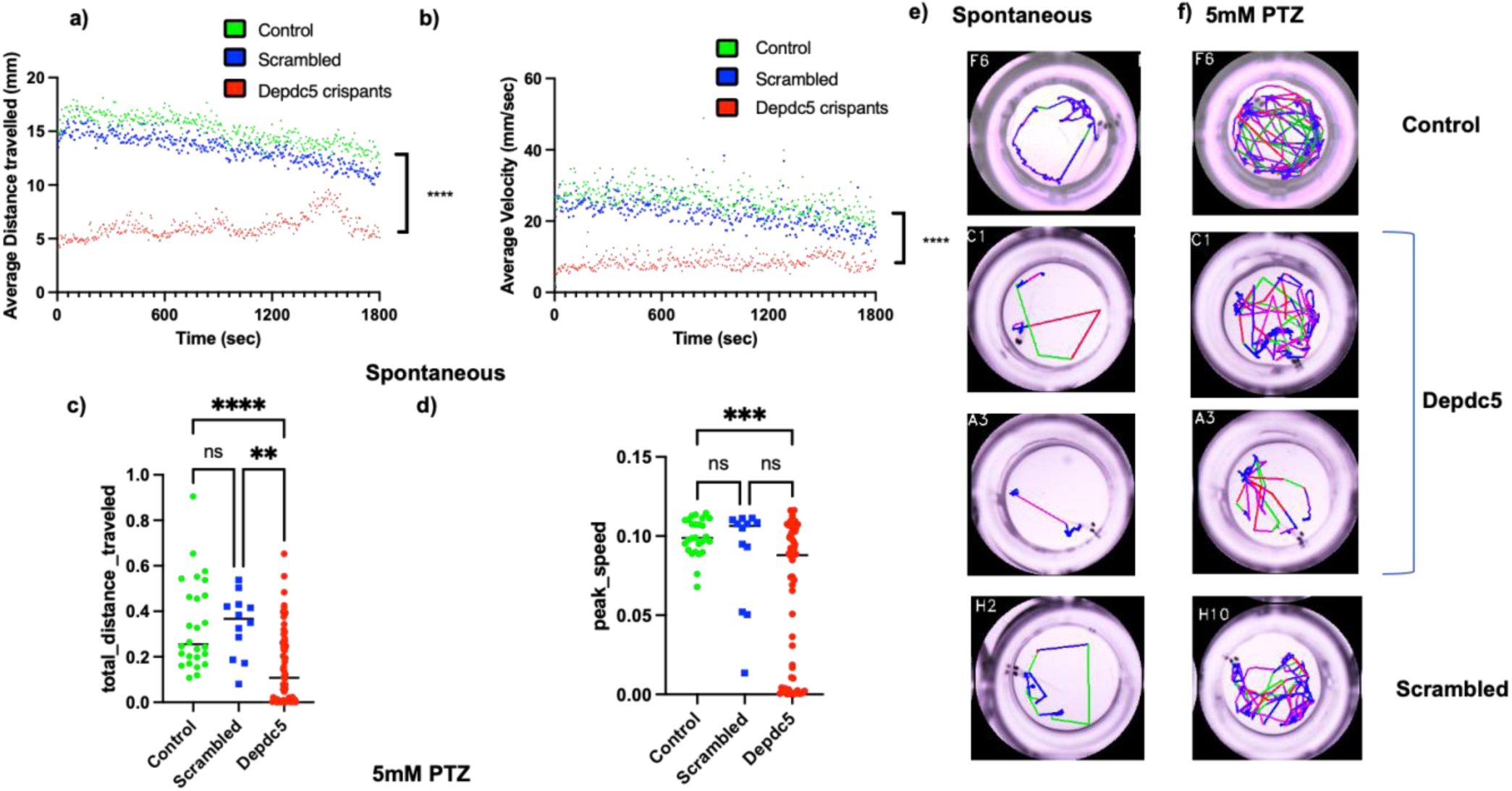
*Depdc5* CRISPants display abnormal spontaneous and PTZ induced swimming behavior. 5 dpf *depdc5* CRISPants and uninjected and scrambled controls were plated in a clear 96 well and the spontaneous swimming behavior was observed in 30-minutes dark cycle, utilizing the Noldus Danio Vision. Representative *depdc5* CRISPants display a) a decrease in the average distance travelled b) and decreased velocity compared to the uninjected and scrambled controls. Further, utilizing the Kestrel MCAM viewer, under the PTZ induced trial, we observed the 5 dpf larvae in 10-minutes dark cycle. *Depdc5* CRISPants display c) abnormal distance travelled while overall being slower compared to the scrambled and un-injected control, d) Peak speed of the *depdc5* CRISPants show decrease as well as a wide range velocity compared to uninjected and scrambled controls, e), f) representative well images of spontaneous *depdc5* CRISPants (c,) and control (d,) larvae in the 96 well plate imaged with Kestrel, a,b). control, π=24, scrambled, n=24, *depdc5* CRISPants n=24.Two-way Anova, , (p<0.0001). c,d) control, n=26, scrambled, n≡12, *depdc5* CRISPants, n=58. (p<0.0001)

In addition to assessing untreated larvae for potential seizure-like swim patterns, we evaluated *depdc5* CRISPants exposed to PTZ. The total distance traveled (Figure 5c,5f) from the center of the well as well as peak velocities (Figure 5d) were substantially decreased in the *depdc5* CRISPants compared to controls. We also observed a range of distance and velocity in different larvae. Along with consistently observed hypoactivity, *depdc5* CRISPants, also show a wide range of peak velocities in PTZ-induced conditions.

### *Depdc5* mosaic CRISPants show neuronal hyperexcitability

To determine whether mosaic *depdc5* loss of function CRISPants display abnormal neuronal network activity, we measured brain activity by recording from the optic tectum at 5 dpf (Figure 6a). We observed recurrent and frequent spontaneous epileptiform events, defined as spikes with high amplitude in the *depdc5* CRISPants brains (Figure 6b) compared to the baseline activity observed in controls (Figure 6c,d). We further separated the larvae into two groups i) non-seizing with baseline activity or no spikes and ii) seizing with high amplitude spikes (> 0.2mV). 53% of surviving *depdc5* CRISPants (heartbeat and blood flow observed for the entirety of a 30-minute trial) demonstrated seizure-like electrical activity (Figure 6e,f) compared to 5% of uninjected and 22% of scrambled injected controls from the same clutch. Our results are consistent with increased neuronal hyperexcitability associated with *depdc5* loss of function.

**Figure 6.**
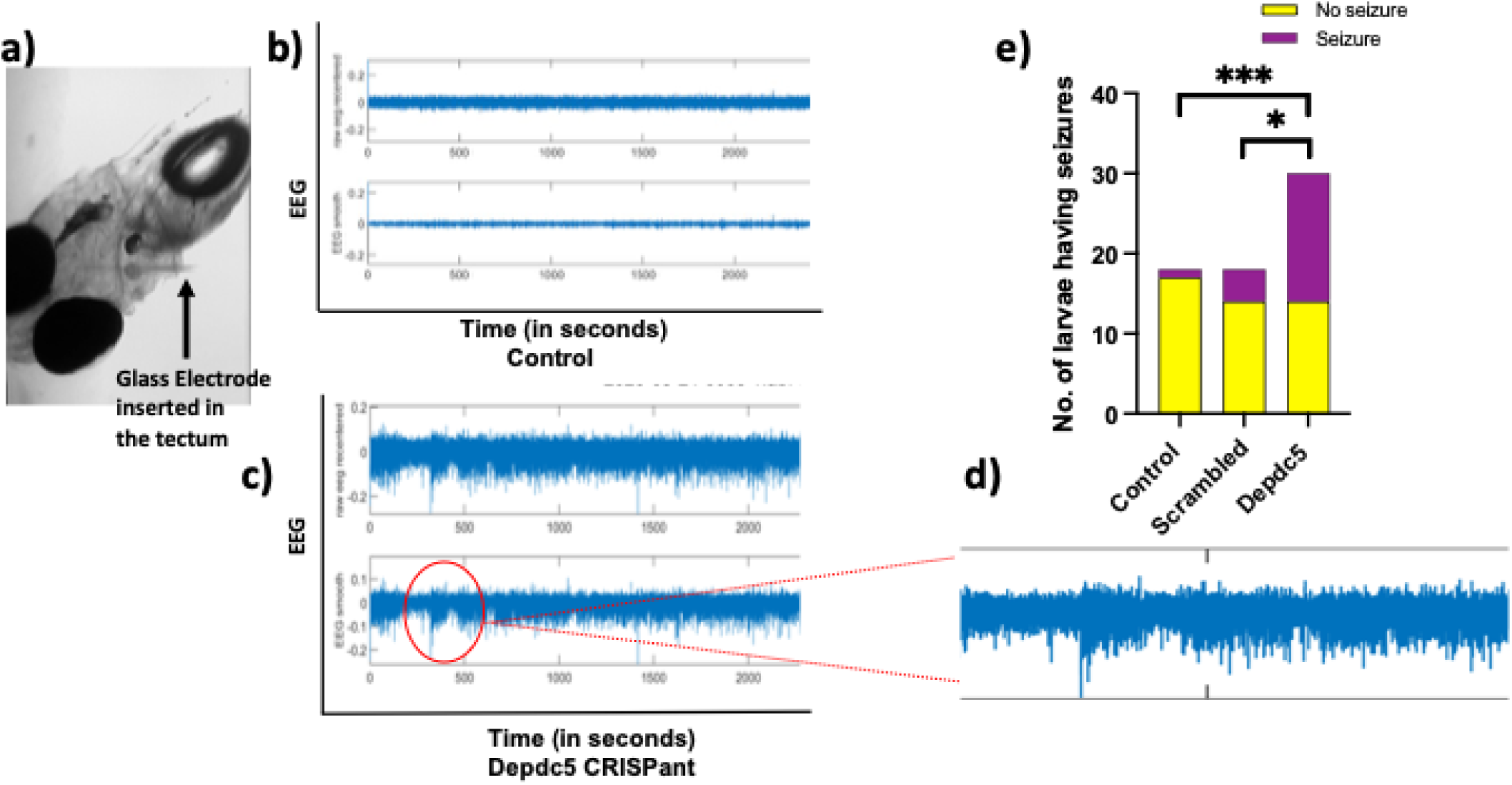
*Depdc5* mosaic CRISPants show hyperexcitability during electrophysiological extracellular recordings from the optic tectum of the larvae. a) saline electrode Inserted carefully in the optic tectum of the larvae (5 dpf). b) representative recording of control, c) representative recording of a *depdc5* CRISPant with seizure like activity; with addition of PTZ towards the end of the run, d) enlarged view of deflections observed in a *depdc5* CRISPant. e) graphical representation of the total number of larvae having seizures vs. no seizures in 30 mins of the readings. Chi-square (and Fisher’s exact test) *depdc5* CRISPants vs. scrambled (p<0.05), *depdc5* CRISPants vs. control (p<0.05). *depdc5* CRISPants n=3O (30 alive), scrambled, n=18, control. n=18

Notably, recording from the CRISPants was sometimes challenging, owing to their abnormal morphology as well as their apparent sensitivity to the acute stress of probe insertion and exposure to the larval paralytic agent alpha-bungarotoxin, resulting in the loss of experimental larvae either in preparatory steps or in the middle of the 30-minute experimental duration. As is typical in such experiments, rare control larvae displayed epileptiform patterns.

### Decreased cell number and evidence of apoptosis in *depdc5* mutant brains

Whole larval staining with AO was performed. We evaluated the tectum region of *depdc5* CRISPants for evidence of apoptosis, as indicated by acridine orange positivity, that can detect early apoptosis. Notably, *depdc5* CRISPants displayed a 3-fold increase in AO clusters in the optic tectum as well as the telencephalon. Interestingly, the uninjected and *scrambled* clutch controls did not display any fluorescent clusters (Figure 7). These data indicate that *depdc5* CRISPants undergo early apoptosis in the brain.

**Figure 7.**
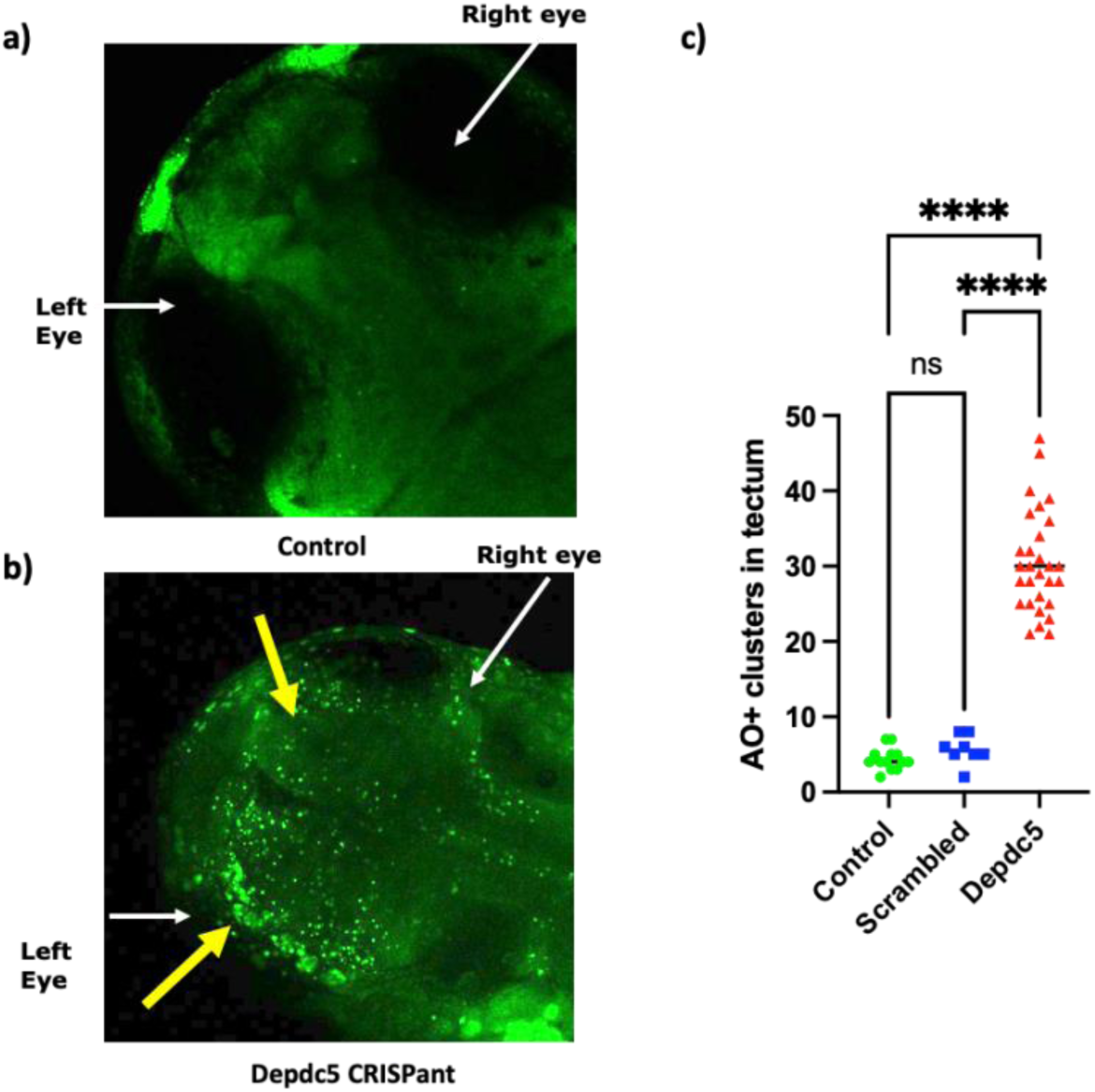
*Depdc5* mosaic CRISPants demonstrate early apoptosis in the larval brain. Live larvae stained with acridine orange that penetrates the early apoptotic cells fluorescing green (yellow arrows), a) representative, control vs, b) *Depdc5* CRISPants. White arrows indicate the left and the right eye of the larvae. Larval age (dpf4). C) graphical representation of the early apoptotic clusters, control, n=13; scrambled, n=8; *depdc5* CRISPants, n=29 One-way Anova, n=29 (p<0.0001)

### *Depdc5* CRISPants display increased posture loss, suggesting seizure-like behavior

Using newly reported methods to assess different types of potentially seizure-like swim patterns,^66^ we observed a significant increase in larval “posture loss” events in *depdc5* CRISPants, particularly in *depdc5*++ CRISPants (Figure 8a) when compared to controls. We did not observe significant differences in the stationary state, normal swim, whirlpool events, or convulsions. In the presence of 5mM PTZ, we observed a significant decrease in stationary behavior, suggesting overall more movement, of *depdc5++* CRISPants compared to controls, as well as the loss of normal swimming behavior in the *depdc5*++ CRISPants. The observed differences in posture loss persisted in the presence of PTZ (Figure 8b) in the *depdc5*++ CRISPants when compared to the *depdc5*+ CRISPants. Further significant increase in posture loss was observed in both *depdc5*++ and *depdc5*+ CRISPants groups when compared to scrambled and uninjected controls. Posture loss events in *depdc5*++ larvae were significantly increased compared to *depdc5*+ larvae. Further, PTZ exposure did not result in changes in normal swim, whirlpool, or convulsion patterns in any group.

**Figure 8.**
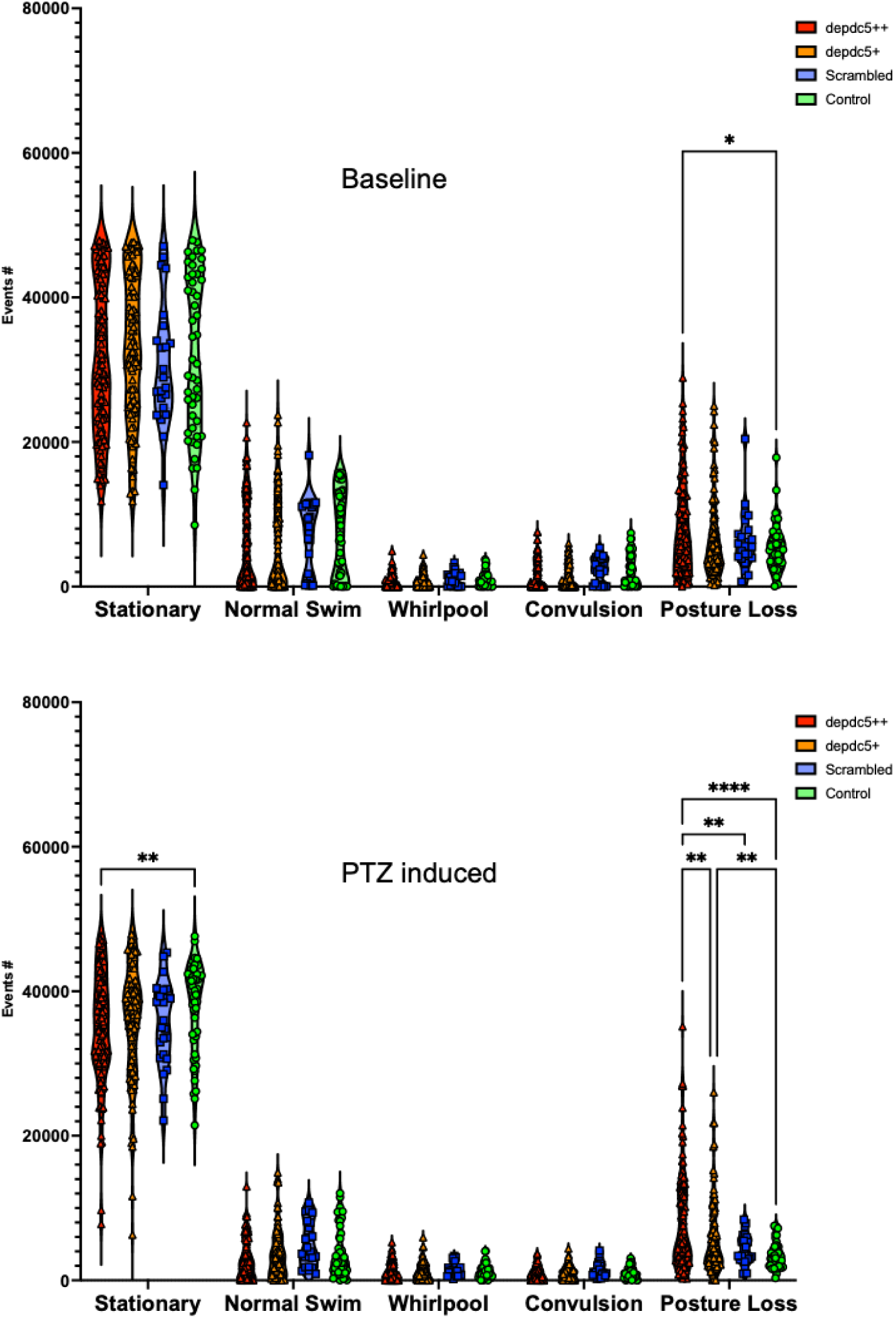
Behavioral ontologies analysis demonstrate changes in posture loss. Freely swimming live larvae of *depdc5* CRISPants *(depdc5++, depdc5+),* uninjected and scrambled controls were recorded for 5-minutes baseline and 5-minutes in 5mM PTZ induced trials, a) The baseline behavior demonstrates significant decrease in posture loss between *depdc5++* and uninjected controls. Additionally, b) in PTZ induced trial a significant difference in stationary as well as decrease in posture loss between the *depdc5++, depdc5+,* uninjected and scrambled controls was observed. Control, n=48; scrambled, n=24; *depdc5* ++ n=113, *depdc5+,* n=9O, Two-way Anova (p<0.0001)

## Discussion

In recent years, the gene *DEPDC5* has been associated with many forms of human epilepsy and sudden death. Loss-of-function variants in *DEPDC5* have been reported in familial and sporadic focal epilepsy disorders,^10, 11^ including mosaic variants associated with focal cortical malformations.^3, 13, 14, 16, 21, 47^ Additionally, our group has reported genetic variants in cases of Sudden Death in Pediatrics (SUDP) and identified potentially contributory variants in *DEPDC5*.^55^ We reported that variants in *DEPDC5* were present in 2 infants who died at 3 months and 5 months of age.^55^ In the current study, we present a mosaic zebrafish larval experimental animal model of *DEPDC5*-related epilepsy. This model recapitulates the classic features observed in individuals with *DEPDC5*-associated epilepsy,^76–78^ including seizure susceptibility and neuronal hyperexcitability as well as early death in a substantial portion of the *depdc5* mutation-positive larvae. *DEPDC5*, a key component of the GATOR-1 complex, is a negative modulator of the mTOR pathway essential for cell growth and development. *DEPDC5* is expressed in neurons in the human brain from the neonatal period through adulthood,^37, 39, 79^ as well as in the nervous systems of vertebrate animals such as mouse and zebrafish.^51^ Zebrafish harbor only one *depdc5* ortholog with strong sequence conservation to humans.^48, 51^ CRISPR-based targeting can be performed without concerns for paralog compensation.

Understanding the importance of *DEPDC5* in human epilepsy and its role in focal epilepsy-related malformations,^14^ we have created mosaic *depdc5* models with demonstrated evidence of mTOR pathway activation indicated by increased presence of PS6 protein. We used constructs that include a fluorescent reporter to label mutant cells. Distinguishing *depdc5* mutant larvae (fluorescence-positive) from non-mutant (fluorescence-negative) larvae that can serve as controls early at the embryonic stage allowed for efficient sorting and assessment of the location (e.g., brain, body) and the level of mosaicism in the regions affected.

Early death has been reported in rodent models of other epilepsy-related genes, such as *Scn1a*^80, 81^ and *Kcna1,*^82^ as well as in *Depdc5* conditional knockout mice.^73–75^ This prompted us to evaluate our *depdc5* CRISPants for premature death. We observed that our *depdc5* mosaic loss-of-function CRISPants demonstrate a high rate of early death at the larval stage compared to scrambled*-*injected and uninjected controls, resembling *DEPDC5*-related SUDP.

In addition to the early death phenotype, we demonstrate that *depdc5* mosaic CRISPants exhibit several features consistent with human mTORopathies.^34^ We demonstrate that *depdc5* CRISPants exhibit abnormal morphology, specifically an elevated head-to-body ratio, potentially consistent with predicted mTOR hyperactivation in the brains of these animals. However, the reduced body length of *depdc5* CRISPants was an unexpected finding and is likely attributable to abnormal, curved body shapes affecting the measurements of mutant larvae.

We evaluated the surviving mosaic *depdc5* mutants for evidence of seizures, with the hypothesis that seizure activity may be related to the early mortality of these models. This issue is especially salient when considering the significance of *DEPDC5* in our cohort if infants and children dying suddenly and unexpectedly, many with neuropathological markers typically found in epilepsy. Previously published metrics reflecting seizure activity include bouts of increased velocity and increased distance traveled.^83, 84^ Surviving *depdc5* CRISPants exhibit abnormal swim patterns, but with *decreased* maximum velocity and distance traveled. These results are consistent with previous studies where hypoactivity was observed in a zebrafish *depdc5* knockout model.^51^ It is possible that morphological abnormalities (e.g., small size, malformed body) or edema in *depdc5* CRISPants limit normal swimming.^43, 59^ Additionally, we observed a range in distance and velocity features in our mosaic *depdc5* mutants in the PTZ induced trials. We also observed increased susceptibility to PTZ in *depdc5* CRISPants compared to uninjected and scrambled controls.

Our electrophysiological data provided evidence that loss of *Depdc5* leads to hyperexcitability, analogous to abnormalities reported in other studies in zebrafish,^48, 51^ rodents,^59, 85^ and human patients.^86^ In local field potential recordings of our mosaic *depdc5* CRISPants, we observed large amplitude events with variable frequencies in different larvae. We observed high amplitude spikes in some larvae that died right after these electrophysiological events, analogous to the terminal seizures seen in some mTOR-related conditional knockout mice.^59, 87, 88^

We investigated possible mechanisms underlying the seizure susceptibility and neuronal hyperexcitability in our mosaic *depdc5* CRISPants, as well as their propensity toward early death. We demonstrated an increase in early apoptosis in clusters of cells, labeled with AO, in *depdc5* CRISPants compared to the uninjected and scrambled controls, suggesting that *depdc5* loss of function leads to early activation of cell death pathways. Because AO predominantly labels cells undergoing membrane alterations typical of early apoptosis, the observed results likely reflect early apoptotic events. Additionally, the apoptotic cells were preferentially located in the larval forebrain and optic tectum, suggesting region-specific vulnerability, as it is reported that telencephalon and mid-brain regions of zebrafish larvae have early *depdc5* expression during development.^48^ The increased AO signal in the *depdc5* CRISPants may provide insights in to the hypothesis that in *depdc5* loss of function associated apoptosis, biological effects including oxidative stress,^89^ disrupt mitochondrial function,^90^ or impair developmental signaling may have a contributory role, that needs to be studied in detail.

Zebrafish larvae exhibit a rich repertoire of motor, sensory, cognitive, and social behaviors that can be systematically described using behavioral ontologies^91–93^. Behavioral ontologies provide standardized descriptors that facilitate integration with genetic and neurobiological data and comparison across studies. An important aspect of *DEPDC5* variants in human epilepsy is that many of them are *de novo* and present in a mosaic state. We sought to correlate the degree of mosaicism, in human studies reflected as variant allele fraction, with abnormal activity. In PTZ exposure trials, *depdc5*++ CRISPants displayed increased and pronounced posture loss events, compared to *depdc5*+ and controls, suggesting that mutational load correlates with phenotypic severity.

### Limitations

We did not observe a phenotype in all *Depdc5* mosaic mutants. Potential reasons for the variability in phenotypes across our models may include variability in which subsets of neurons carried the variant, as well as variability in variant allele fraction from animal to animal. One explanation for lack of behavioral seizures may be that our relatively slow frame rate does not allow for detection of short bouts of fast swimming. We also may have underestimated the abnormal electrophysiological activity of the mutants given the relatively short experimental window (30 minutes). Longer recordings were not possible because mutant larvae were so highly vulnerable to dying during recording compared to controls. It is also possible that recording behavioral swim patterns and LFPs in younger larvae (2-4 dpf) may reveal more abundant and frequent abnormalities since *depdc5* expression is higher at early embryonic ages. Additionally, AO staining does not distinguish between apoptosis and necrotic cells with compromised membrane integrity. Complementary assays to AO, e.g., Caspase-3 immunostaining, would provide further support for the hypothesis that there is early apoptosis in the CRISPants.

## Conclusions

In summary, our *depdc5* mosaic CRISPants demonstrate key features of human *DEPDC5*-related epilepsy, including premature death and, in surviving CRISPants, evidence of seizure susceptibility to PTZ, and hyperexcitability that are more striking in animals with generally high numbers of mutant cells. While preliminary, these findings also support the role of *DEPDC5* in early child mortality, as induced from cases without a history of seizures in which deleterious variants were identified. Mosaic zebrafish models offer a key advantage over traditional germline (non-mosaic) models, particularly for studying genes like *DEPDC5* that are involved in mosaic focal epilepsies. A standard (germline) knockout, in which all cells lack the gene of interest, may lead to lethality or severe phenotypes that do not mimic the focal nature of many human epilepsies. Mosaic models more accurately reflect the somatic, postzygotic mutations observed in patients with FCD and other mTOR - related epilepsies, where only a subset of brain cells carries the pathogenic mutation. The juxtaposition of mutation-positive with mutation-negative cells in mosaic whole-organism models will allow future studies to examine the interaction of the mutant vs. WT cells, which is critical for understanding cell-autonomous vs. non-cell-autonomous mechanisms of mTOR pathway disruption and subsequent seizure generation. Future models in which only a subset of neurons are mutation-positive can be developed with neuronal-specific Cre drivers to isolate which neuronal populations are important for normal *Depdc5* activity. All of these models provide a system in which future screening of drugs for children and adults with *DEPDC5*-related mosaic epilepsies can be evaluated.

## Author contributions

Experiments were designed by ST and AP with input from the entire research team. ST and HYK designed the UFlip construct. ST conducted the experiments, with CML conducting the electrophysiology experiments. Data analysis was conducted by ST, CML, JP, PC, and AP. AP reviewed all experimental data and analyses. ST, RDG, and AP provided funding for the study. ST and AP drafted the manuscript. All the authors read and approved the final manuscript.

## Acknowledgments

The authors thank the dedicated staff of the Boston Children’s Hospital Aquatic Resources Program (ARP). We would like to thank the BCH Cellular Imaging Core; RRID:SCR_026485, funded by NIH P50 HD105351. We are grateful to Dr. Alexander Rotenberg and the Experimental Neurophysiology Core (ENC) at BCH. We thank the Ramona Kestrel team for their technical support and the MCAM software. We are grateful to our former lab manager Dr. Laura Turner for her support.

