## Supplement Figures 1 & 2 for "Early death and neuronal abnormalities in *depdc5* loss-of-function mosaic zebrafish models"

**Supporting Data**

**Figure S1**

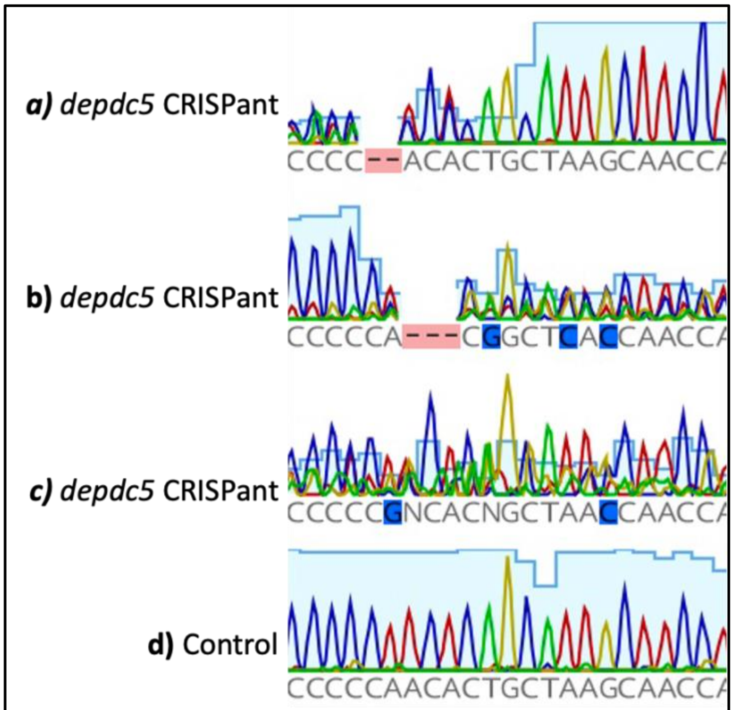


**Figure S1. Sanger sequencing shows targeting *depdc5*.** The Sanger sequencing graphs showing the successful targeting of *depdc5* in the *depdc5* CRISPants (a,b,c) compared to the (d) control larvae.

**Figure S2**

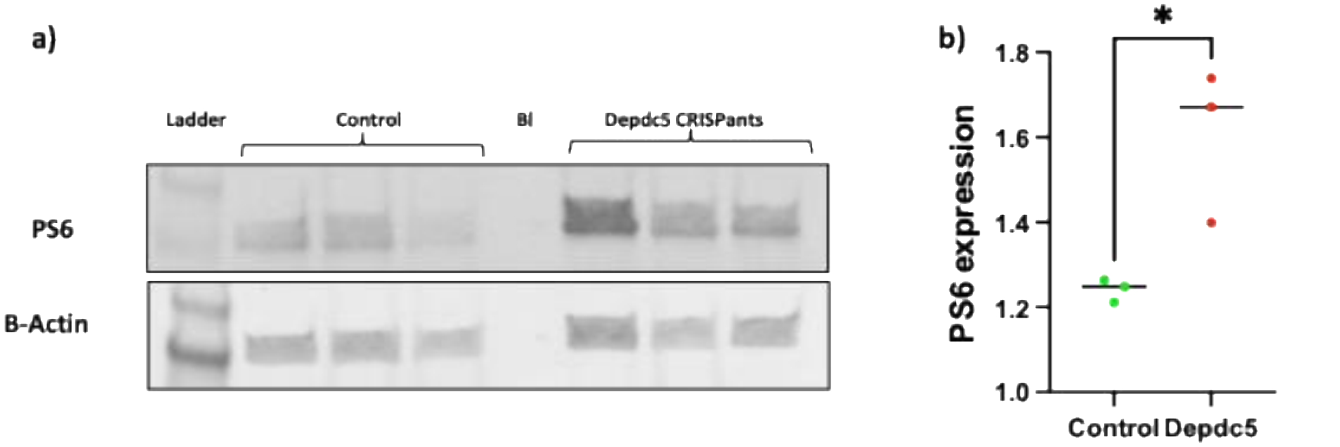


**Figure S2. Overexpression of PS6 in the *depdc5* CRISPants** a) PS6 protein expression in control larval protein compared to protein from *depdc5* CRISPants, demonstrating Phospho-S6 activity in the *depdc5* CRISPants. Housekeeping protein Beta-actin expression in the lower row, for normalization. b) Graphical representation of the protein expression. Control, n=3; Depdc5 CRISPants, n=3 (each sample is a pool of 5 dpf5 larval protein)
